# Stabilization of RRBP1 mRNA via an m6A-dependent manner in prostate cancer constitutes a therapeutic vulnerability amenable to small-peptide inhibition of METTL3

**DOI:** 10.1101/2023.09.04.556177

**Authors:** Yuqing Feng, Zenghui Li, Jingwei Zhu, Cheng Zou, Yu Tian, Jiangling Xiong, Qinju He, Wenjun Li, Hao Xu, Bin Xu, Junfeng Shi, Dingxiao Zhang

## Abstract

Mounting evidence has implicated the RNA m6A methylation catalyzed by METTL3 in a wide range of physiological and pathological processes, including tumorigenesis. The detailed m6A landscape and molecular mechanism of METTL3 in prostate cancer (PCa) remains ill-defined. We find that METTL3 is overexpressed in PCa and correlates with worse patient survival. Functional studies establish METTL3 as an oncoprotein dependent on its m6A enzymatic activity in both AR+ and AR- PCa cells. To dissect the regulatory network of m6A pathway in PCa, we map the m6A landscape in clinical tumor samples using m6A-seq and identify genome-wide METTL3-binding transcripts via RIP-seq. Mechanistically, we discover RRBP1 as a direct METTL3 target in which METTL3 stabilizes *RRBP1* mRNA in an m6A-dependent manner. RRBP1 positively correlates with METTL3 expression in PCa cohorts and exerts an oncogenic role in aggressive PCa cells. Leveraging the 3D structural protein-protein interaction between METTL3 and METTL14, we successfully develop two potential METTL3 peptide inhibitors (RM3 and RSM3) that effectively suppress cancer cell proliferation *in vitro* and tumor growth *in vivo*. Collectively, our study reveals a novel METTL3/m6A/RRBP1 axis in enhancing aggressive traits of PCa, which can be therapeutically targeted by small-peptide METTL3 antagonists.

## INTRODUCTION

Prostate cancer (PCa) is a major cause of cancer-related deaths in men globally [1]. The treatment options are currently limited for advanced PCa patients, and androgen deprivation therapy (ADT) is the mainstay of therapy. Unfortunately, ADT usually fails to endow a durable response and most patients eventually develop castration-resistant PCa (CRPC), which is currently incurable. Therefore, development of new therapeutics for aggressive PCa variants, especially CRPC, is urgently needed. So far, our understanding of PCa progression is incomplete, warranting further detailed mechanistic studies.

N6-Methyladenosine (m6A) is the most prevalent internal modification for eukaryotic RNA and influences nearly every stage of RNA metabolism, including transcription, splicing, decay, export and translation [2, 3]. As a dynamic and reversible process, m6A modification is decorated by Writer complex (i.e., methyltransferase including mainly METTL3, METTL14 and WTAP), removed by Erasers (i.e., demethylases FTO and ALKBH5) and recognized by Readers (i.e., m6A-binding proteins including YTHDF1/2/3, IGF2BP1/2/3, HNRNPA2B1 and others) [2]. Increasing evidence has firmly implicated m6A pathway in a variety of cancer development, but in a context-dependent manner. For example, METTL3 has been reported to function as an oncogenic factor in glioblastoma [4], colorectal carcinoma [5], leukemia [6], and bladder cancer [7]. Whereas, a tumor suppressive role for METTL3 has also been suggested in ocular melanoma [8] and triple-negative breast cancer [9]. Consequently, METTL3 has been recently emerged as an attractive target for anticancer drug development.

Conceptually, bioinformatic analyses have suggested an important role for METTL3-mediated m6A pathway in prostate tumorigenesis [10-13]. Experimentally, there are few papers reported the biological role of METTL3 in PCa cells and identified few m6A targets on an individual gene basis such as GLI1[14], MYC[15], KIF3C[16], ITGB1[17], USP4[18]. However, several key questions remain unanswered: **1**) a landscape of m6A methylation on PCa transcriptome in clinical tumors is uncharacterized; **2**) the genome-wide targets of METTL3 in PCa cells remain elusive; **3**) evidence supporting METTL3 as a valid therapeutic target for treating PCa lacks, in that previous studies mainly used genetic knockdown approaches instead of therapeutic interference with m6A signaling modulators. Here, we systematically investigate METTL3 expression, functions, and its direct targets in PCa. We find that METTL3 is oncogenic in human PCa by methylating and thus stabilizing *RRBP1* mRNA in an m6A-dependent manner. Using m6A-seq and RIP-seq in clinical samples, we further reveal a gene regulatory network governed by METTL3-mediated m6A signaling. Importantly, although a small-molecule METTL3 inhibitor STM2457 has been recently introduced for treating blood cancers [6], its efficacy against solid PCa is less effective. Alternatively, we develop for the first-time peptide inhibitors (linear RM3 and stapled form RSM3) that drastically inhibit METTL3 enzymatic activity and induce METTL3 degradation, leading to a strong anticancer toxicity in PCa both in vitro and *in vivo*.

## RESULTS

### 1. Upregulation of METTL3 in PCa correlates with poor survival

Recently, we have comprehensively dissected, genomically and transcriptomically, the m6A pathway (24 genes) as a whole in PCa using TCGA cohort [19]. We found that many of the m6A pathway genes were upregulated, while a few genes (including eraser FTO) downregulated, in primary PCa (pri-PCa) versus (vs.) normal tissues [19], indicating an overactivation of m6A pathway. To further investigate the potential roles of m6A modification in PCa evolution in detail, we focused on METTL3 as it’s the only enzymatic subunit in the Writer complex. Analysis of *METTL3* mRNA levels in Oncomine database revealed that *METTL3* was overexpressed in pri-PCa vs. normal/benign prostate tissues (Fig. 1A). This result was further confirmed by analyzing both the curated [20] and noncurated (i.e., provisional) TCGA cohorts (Fig. 1B). Interestingly, examination of large clinical RNA-seq datasets indicated a further increase of *METTL3* levels in CRPC va. pri-PCa (Fig. 1C). In an attempt to understand the molecular basis underpinning *METTL3*’s dysregulation in PCa, we surveyed its mutational landscape. Although we did see an increase in METTL3 alteration in CRPC vs. pri-PCa cohorts, it was in total mutated at a very low frequency (Fig. S1A), suggesting that its mis-expression was not due to genomic alterations. Clinically, a strong positive correlation between *METTL3* expression and tumor Gleason Score (GS) was observed (Fig. 1D), consistent with a gradual upregulation of *METTL3* in GS high (>7) vs. low (≤7) tumors (Fig. 1B). Also, the *METTL3* mRNA levels adversely associated with PCa patients’ overall survival in multiple datasets (Fig. 1E). Experimentally, we determined METTL3 protein expression in a panel of prostate normal and cancerous cell lines, finding a general overexpression in PCa cells vs. immortalized normal prostatic epithelial cells RWPE1 (Fig. S1B). Altogether, our data established a potential oncogenic role of METTL3 in PCa initiation and castration-resistant progression.

**Fig 1.**
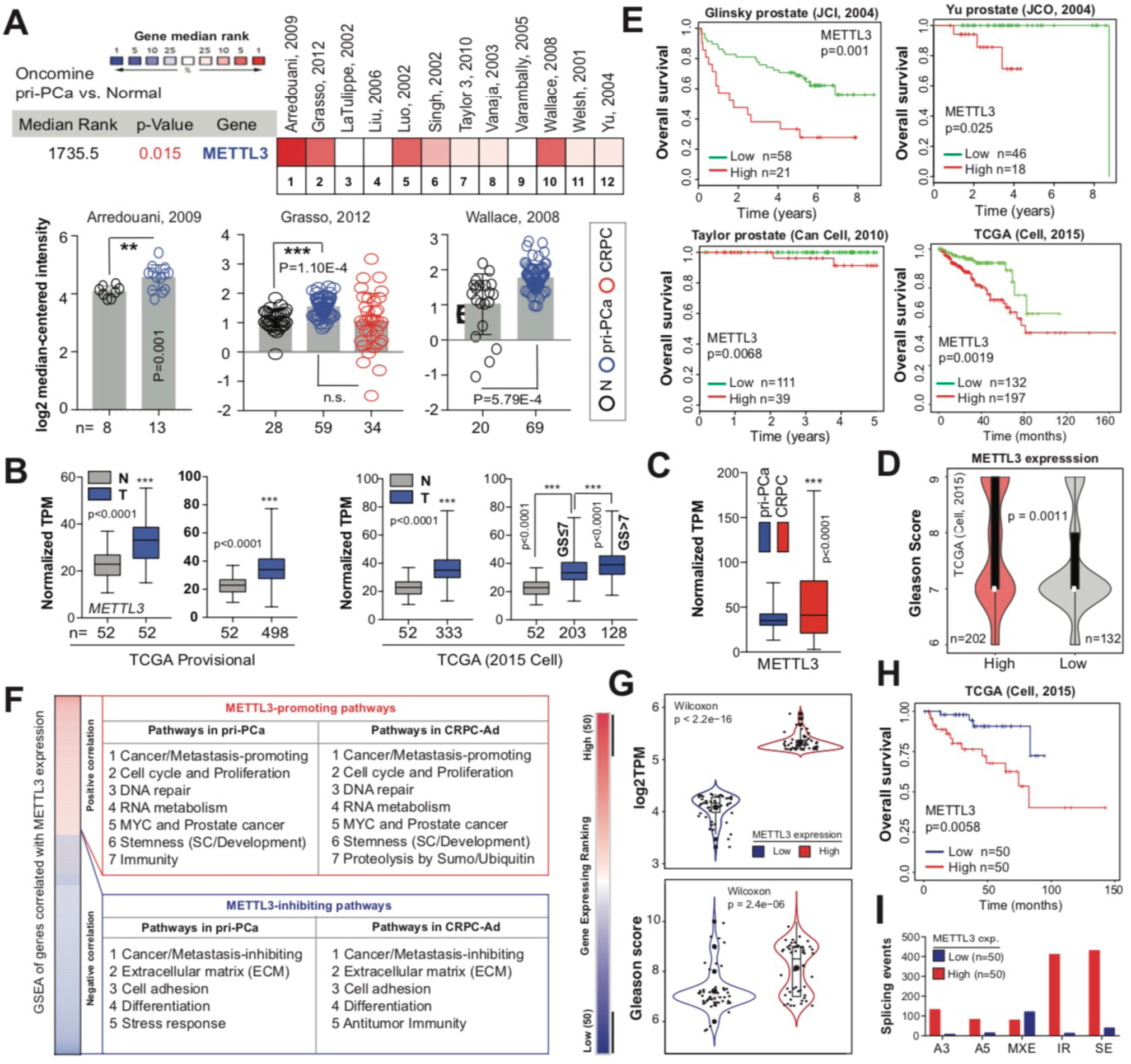
Increased METTL3 expression in PCa correlates with multiple oncogenic pathways and worse patient survival. **A.** Oncomine analysis showing increased *METTL3* expression in PCa vs. normal or benign tissues. The bottom illustrates the details in three representative datasets. **B.** Overexpression of *METTL3* at mRNA level in both curated (2015 Cell) and noncurated pan-cancer PCa TCGA cohorts. **C.** Comparison of *METTL3* expression reveals an upregulation in CRPC vs. pri-PCa samples. **D** and **E.** High *METTL3* mRNA levels correlate with increase GS (D) and worse patient overall survival (E) in indicated datasets. Survival *p*-value was determined using the Log-Rank test. **F.** GSEA of genes co-expressed with *METTL3* in curated pri-PCa (TCGA) and CRPC (SU2C-PCF) cohorts. **G-I**. Fractionation of TCGA pri-PCa cohort (2015 Cell) into *METTL3* high and low groups, and comparison of *METTL3* expression (**G**, up), GS (**G**, bottom), patient’s survival outcome (**H**), and splicing landscape (**I**) showing *METTL3* high group being more aggressive.

### 2. A protumorigenic role of METTL3 is associated with multiple cancer-related pathways

Many roles of METTL3-mediated m6A signaling have been proposed in development and cancer [2, 3]. To comprehensively dissect the molecular functions of METTL3 in PCa, we performed a gene-coexpression analysis, coupled with gene set enrichment analysis (GSEA), to identify biological gene signatures and pathways that may be regulated by METTL3. Both curated TCGA pri-PCa [20] and CRPC [21] cohorts were examined and similar repertoires of biological pathways were identified (Fig. 1F and Table S1), indicating conserved roles of METTL3 in both treatment-naïve and treatment-resistant PCa. Globally, METTL3 positively regulated proliferation (evidenced by enrichment of pathways tied to cell cycle progression and DNA repair), MYC and PCa-related signatures, RNA metabolism (such as transcription and splicing), and stemness to eventually promote cancer development and progression (Fig. 1F and Fig. S1C). Supporting our analyses, the involvement of m6A pathway in all these processes have been documented in diverse biological contexts [3, 22, 23]. For example, it has been reported that METTL3 plays an oncogenic role in multiple cancer types [22] by regulating tumor stemness [24] and RNA stability and splicing [25, 26], among other mechanisms. Conversely, genes negatively correlated with *METTL3* expression were enriched in pathways associated with extracellular matrix (ECM) and cell adhesion, differentiation, stress response and/or anti-tumor immunity (Fig. 1F and Table S1), indicating a cancer/metastasis-inhibiting function. To further corroborate these results, we fractionated TCGA pri-PCa cohort into two extremes (Fig. 1G) and asked whether high or low *METTL3* levels in PCa was indeed associated with tumor aggressiveness. Expectedly, tumors expressing highly (vs. lowly) the *METTL3* exhibited elevated GS (Fig. 1G, bottom), worse survival outcome (Fig. 1H), and distinct splicing landscapes (Fig. 1I). The total dysregulated splicing events (5.77-fold) and intron retention (IR; 31.62-fold) were specifically upregulated in the *METTL3* high group (Fig. 1I). We have recently shown that the severity of RNA splicing disruption correlates with increased aggressiveness [27]. Collectively, higher *METTL3* expression predicted a molecularly aggressive phenotype in PCa, consistent with recent reports [15, 18].

### 3. Attenuation of METTL3 inhibits PCa progression *in vitro* and *in vivo*

A critical role of METTL3 in mobility of AR- CRPC cells (i.e., DU145 and PC3) has been reported recently [18]. To comprehensively dissect the roles of METTL3-mediated m6A pathway in both AR+ and AR- cells, we knocked down METTL3 in LNCaP and PC3 cells. Two independent shRNA-mediated depletion of METTL3 (Fig. 2A and Fig. S2A) significantly reduced the total m6A levels in PCa cells (Fig. 2B), and concurrently led to a remarkable impairment of cell growth in these lines, as evidenced by the colony formation assay (Fig. 2C). Moreover, trans-well (Fig. 2D) and wound-healing (Fig. 2E) assays revealed that attenuation of *METTL3* significantly decreased cell mobility in PC3 and DU145 cells, with PC3 being the most affected one. Quantitation of apoptosis via Annexin V staining indicated that loss of *METTL3* caused a significant increase in apoptosis (Fig. 2F and Fig. S2B) in both AR+ LNCaP and AR- lines, validating the reduced proliferation phenotype (Fig. 2C). Moreover, knockdown (KD) of METTL3 significantly inhibited PCa sphere formation (Fig. 2G), as well as the sphere size (Fig. S2C), indicating a requirement of METTL3 for optimal PCa stemness. Alternatively, we obtained similar results by depleting *METTL3* with two independent siRNAs in PCa cells. Briefly, siRNA treatment significantly reduced METTL3 expression both at mRNA (Fig. S2D) and protein (Fig. S2E) levels, leading to a global decrease in RNA m6A methylation (Fig. S2F), clonal development capacity (Fig. S2G), viability (Fig. S2H), and migration (Fig. S2I) in both DU145 and PC3 cells. All these abovementioned *in vitro* assays strongly established a protumorigenic role of METTL3 in PCa. To next demonstrate the role of METTL3 *in vivo*, we generated subcutaneous tumors in BALB/c athymic nu^-^/nu^-^ (nude) male mice implanted with PCa cells stably expressing control or *METTL3*-targeting shRNAs. As shown in Fig. 2H, *METTL3* KD inhibited the growth of both PC3 and DU145 xenograft models. Collectively, these data convincedly demonstrated that METTL3 is oncogenic in PCa development and progression.

**Fig 2.**
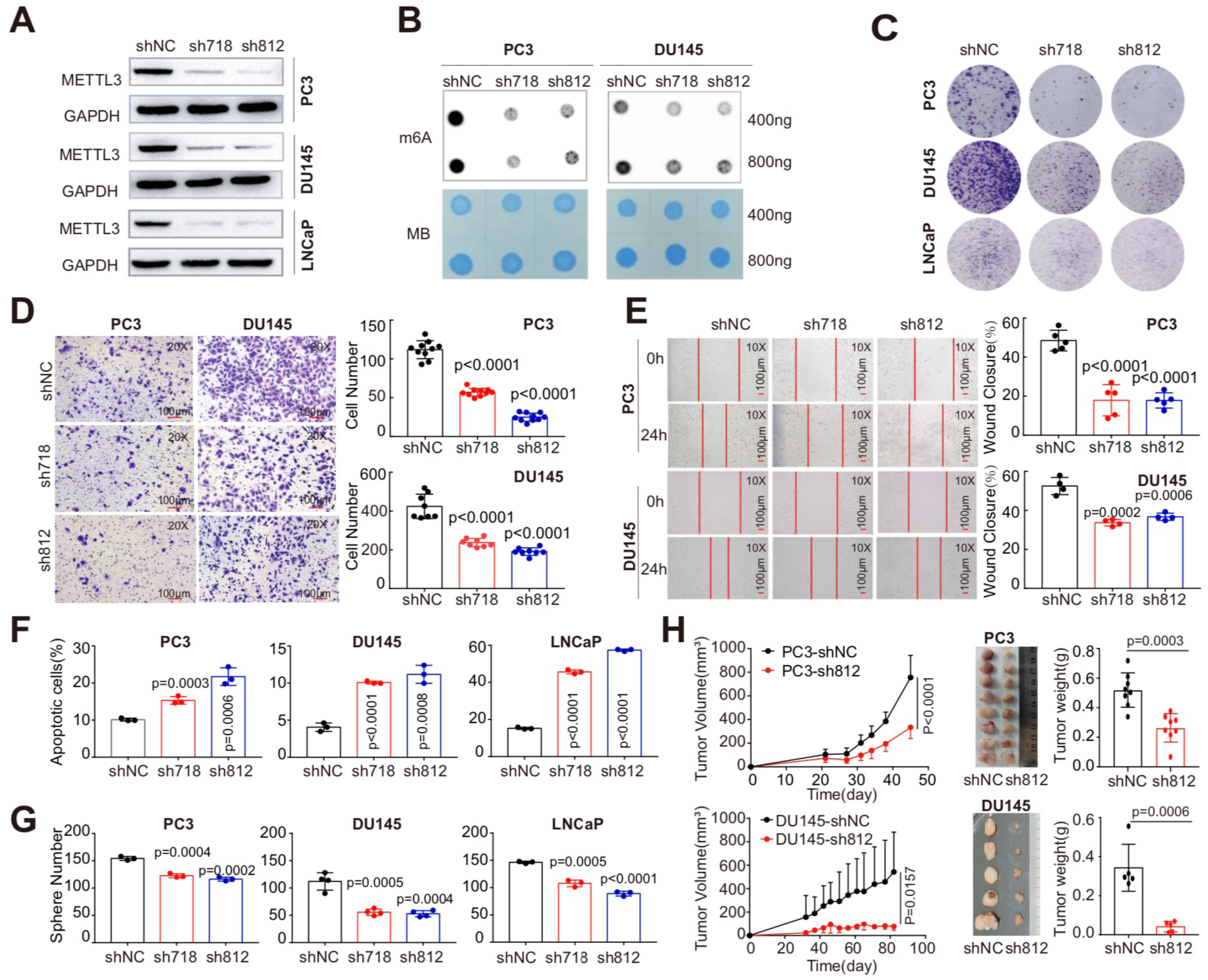
Knock-down of METTL3 significantly inhibits PCa progression *in vitro* and *in vivo*. **A.** Western blot analysis showing METTL3 abundance in three PCa cell lines transduced with scrambled (shNC) or *METTL3*-specific hairpin shRNAs (sh718 and sh812) and probed with indicated antibodies. GAPDH served as a loading control. **B.** Dot blot assay showing the global m6A levels in indicated cells treated with different shRNAs. Methylene blue (MB) staining served as a loading control. **C.** Knocking down *METTL3* inhibits clonal development in indicated PCa cells. **D** and **E.** Cell mobility evaluated by Trans-well (D) and wound-healing (E) assays showing that *METTL3* knock-down inhibits cancer cell migration. **F.** Flow cytometry analysis showing an increased cellular apoptosis in indicated PCa cell lines upon *METTL3* depletion. **G.** Knocking down *METTL3* inhibits sphere formation in both AR+ and AR- PCa cell lines. **H.** Depletion of *METTL3* inhibits the growth of PC3 and DU145 xenograft tumors *in vivo* (n=5 for each group). Shown are the tumor growth curves (left), endpoint tumor images (middle), and tumor weight (right) of indicated PCa models.

### 4. METTL3 promotes prostate tumorigenesis in an m6A-dependent manner

METTL3 forms an obligate heterodimer with METTL14 to exert its methyltransferase activity [2]. To determine whether METTL3’s function of accelerating PCa aggressiveness was dependent on its m6A catalytic activity, we successfully established stable cell lines expressing wild-type (WT) METTL3 or its catalytic-dead mutant (aa395-398, DPPW→APPA). A lack of methyltransferase activity of this mutant has been previously described [28, 29]. Overexpression of the WT, but not the mutant, METTL3 (Fig. 3A) resulted in a significant increase in global m6A levels in both DU145 and PC3 cells by dot-blot assay (Fig. 3B), which was further confirmed by quantification of RNA methylation status via EpiQuik m6A Quantification Kit (Fig. 3C). A series of cell-based assays in two different PCa cell lines showed that only transduction of WT (compared to the mock or mutant-expressing) METTL3 lentivirus further promoted cell clonal development (Fig. 3D), proliferation (MTT assay; Fig. 3E), and migration ability (trans-well assay; Fig. 3F). Importantly, xenograft tumor assay indicated that the catalytic-dead METTL3 (vs. WT) failed to promote DU145 tumor development *in vivo*, as evidenced by indistinguishable difference in tumor growth dynamics (Fig. 3G) and weight (Fig. 3H) between mutant and control groups. Collectively, these results strongly indicated that the m6A catalytic activity of METTL3 is required for its role in advancing prostate tumorigenesis.

**Fig 3.**
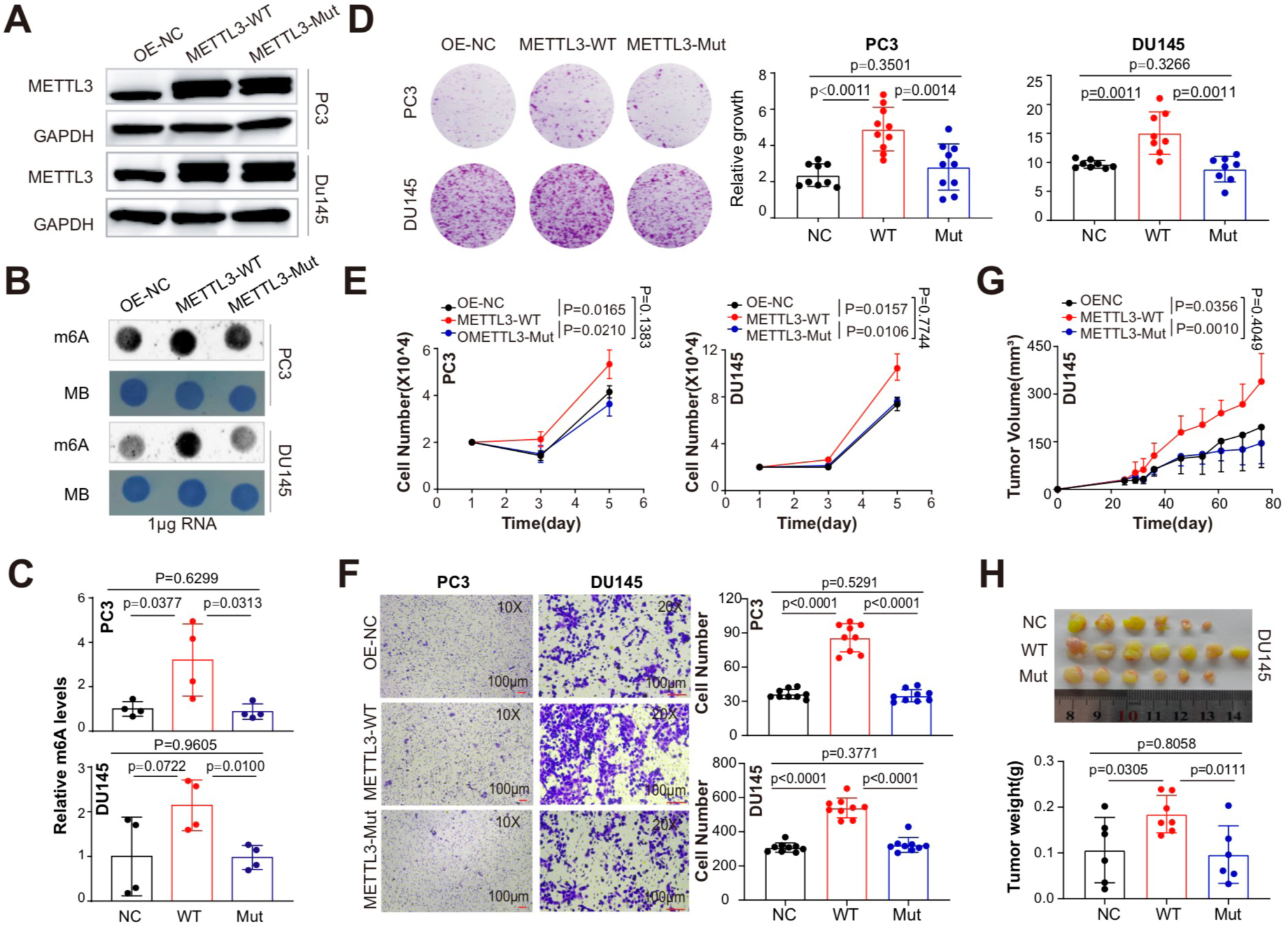
METTL3 overexpression promotes PCa progression in a m6A-dependent way. **A.** Western blot analysis showing overexpression of WT or catalytically dead (i.e., mutant) METTL3 in PCa cells. GAPDH served as a loading control. **B.** Dot blot assay showing the global m6A levels in indicated cells treated with different overexpressing constructs. Methylene blue (MB) staining served as a loading control. **C.** Relative m6A levels in indicated conditions detected via EpiQuik m6A quantification kit. **D-F.** Overexpression of WT, but not the mutant, METTL3 promotes CRPC clonal development (**D**), proliferation (**E**), and Trans-well migration (**F**) in both DU145 and PC3 cells *in vitro*. **G** and **H.** The m6A enzymatic activity is required for METTL3 in enhancing PCa development *in vivo*. Shown are the tumor growth curves (**G**), endpoint tumor images and tumor weight (**H**) of DU145 model.

### 5. Identification of global METTL3 targets in PCa cells

To directly unravel the mechanisms of action of METTL3 in PCa cells, we utilized a multi-omics approach. We first performed RNA-seq analysis in PC3 cells with or without *METTL3*-KD, finding a set of 177 differentially expressed genes (DEGs; 66 up- and 111 down-regulated) (Fig. 4A and Table S2). Gene ontology (GO) analysis indicated that genes upregulated in *METTL3*-KD cells were enriched in pathways tied to inflammation/immunity and cell death (Fig. 4B), whereas genes downregulated were enriched in pathways associated with migration/adhesion, differentiation and proliferation, in line with an oncogenic role of METTL3. The m6A writer complex binds RNA to decorate m6A modification [23]. Next, we performed RNA immunoprecipitation sequencing (RIP-seq) experiments in both AR+ LNCaP and AR- PC3 cells with an METTL3-specific antibody. As shown in Fig. 4C, 2,738 and 1,382 potential METTL3 target genes were identified in PC3 and LNCaP, respectively, with 849 genes being shared between these two cell lines (Fig. 4D and Table S3). Consistently, GO analysis of genes bound by METTL3 revealed, overall, a similar pattern of biological pathways enriched in LNCaP and PC3 cells (Fig. 4E), highlighting that METTL3 may play similar functional roles in both AR+ and AR- cells. To further identify genes that were directly regulated by METTL3 in PCa cells, we integrated the RNA-seq and RIP-seq data by overlapping DEGs and genes bound by METTL3 protein (Fig. 4D), and identified 11 key genes (Fig. 4F). Among them, SYNE2 [30] and HEG1 [31] have been reported to be regulated by m6A modification in different context, indicating validity of our data. We chose RRBP1 for further investigation as it showed the most significant elevation in mRNA expression upon *METTL3*-KD.

**Fig 4.**
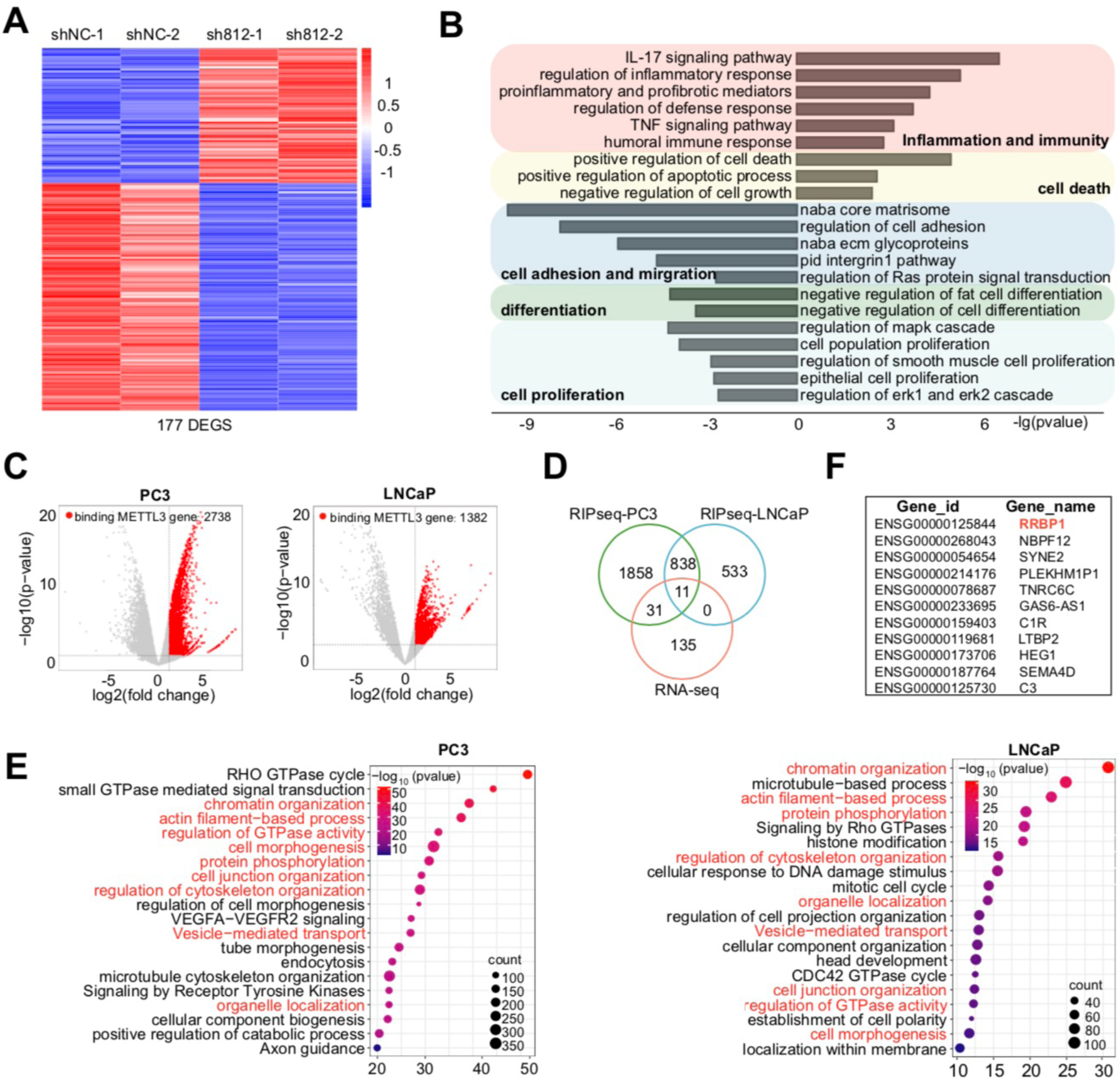
Identification of the genome-wide METTL3 targets in PCa cells. **A.** Heatmap of differentially expressed genes (DEGs) identified in PC3 cells by RNA-seq (fold change (FC)≥1.5 and FDR<0.1). **B.** GO analysis of DEGs as shown in A. **C.** Volcano plot of METTL3-bound genes identified in AR+ LNCaP and AR- PC3 cells by RIP-seq (FC≥2 and FDR<0.1). **D.** Overlap between DEGs and METTL3-bound genes in indicated contexts. **E.** GO analysis of METTL3-bound genes showing that a significant proportion of enriched pathways are commonly enriched in both LNCaP and PC3 cells. **F.** List of the 11 overlapped genes from D.

### 6. METTL3 accelerates PCa progression through upregulating oncogenic RRBP1

In line with an upregulation of METTL3 in pri-PCa and CRPC, similar pattern was found for *RRBP1* mRNA levels (Fig. 5A), implying a positive relationship between them. Quantitative RT-PCR (qPCR) confirmed a reduction in RRBP1 expression at both mRNA (Fig. 5B) and protein (Fig. 5C) levels upon *METTL3*-KD. Notably, the enzymatic activity of METTL3 was required for this phenotype, as only the exogenous expression of WT, but not the mutant, METTL3 strongly upregulated RRBP1 protein expression (Fig. 5D). To further demonstrate *RRBP1* as a direct substrate bound by METTL3, we performed RIP-qPCR to reveal that METTL3 strongly bound *RRBP1* transcripts in both DU145 and PC3 cells (Fig. 5E) and the m6A modification on *RRBP1* mRNA by m6A-qPCR was significantly reduced upon *METTL3*-KD (Fig. 5F). Analysis of *RRBP1* sequences and METTL3 RIP-seq peaks unraveled multiple consensus m6A motifs (RRACH) in its 3’-UTR region (Fig. 5G). Mutagenesis assay was performed with luciferase (Luc) reporters containing either a WT or a mutated (MUT) 3’-UTR placed after the coding region of a firefly luciferase. Results indicated that the luciferase activity of WT, but not the MUT, reporter was significantly reduced in DU145 cells when METTL3 was knocked down (vs. shNC control) (Fig. 5H), suggesting a positive regulation of m6A on *RRBP1* mRNA. To further dissect the fate of *RRBP1* mRNA upon m6A methylation, we determined its mRNA stability after transcriptional inhibition with Actinomycin D. Attenuation of METTL3 markedly accelerated *RRBP1* mRNA decay in both DU145 and PC3 cells (Fig. 5I), suggesting that METTL3 upregulated *RRBP1* through mRNA stability in a m6A-dependent manner. Altogether, these results established *RRBP1* as a direct target of METTL3 in PCa cells.

**Fig 5.**
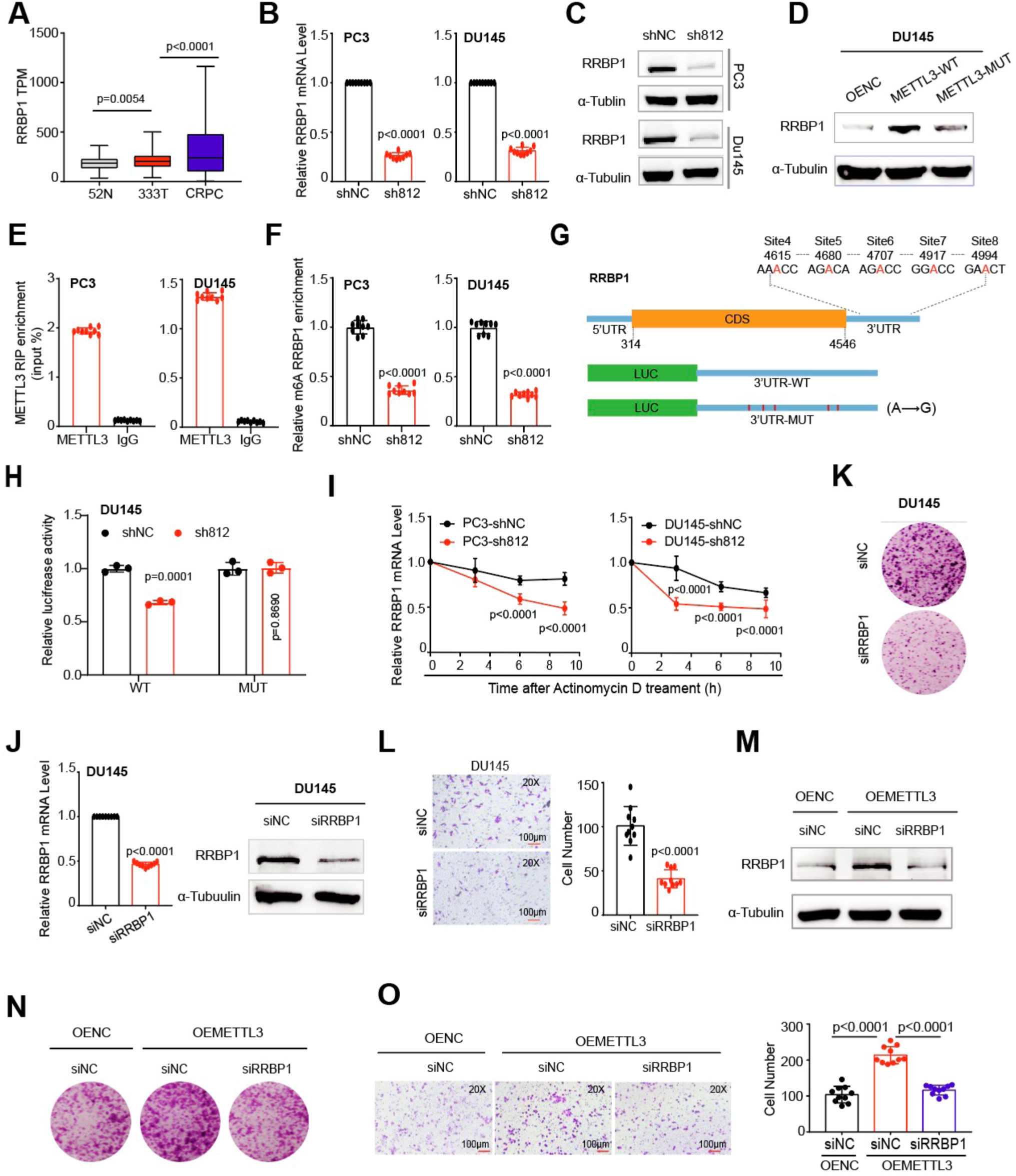
Oncogenic RRBP1 is a direct and functional target of METTL3-mediated m6A signaling in promoting PCa aggressiveness. **A.** Overexpression of *RRBP1* at mRNA level during PCa progression, as analyzed in TCGA (pri-PCa vs. N) and CRPC (vs. pri-PCa) cohorts. **B.** qPCR analysis showing a reduced *RRBP1* expression in PC3 and DU145 cells upon *METTL3* depletion. **C** and **D.** Western blot analysis of RRBP1 protein levels in indicated PCa cells with *METTL3*-KD (**C**) or METTL3 overexpression (**D**). Tubulin served as a loading control. **E.** RIP-qPCR analysis showing an enrichment of METTL3 binding at *RRBP1* transcripts in PCa cells. **F.** m6A-qPCR analysis showing much reduced m6A modifications in *RRBP1* transcripts in PCa cells upon *METTL3*-KD. **G.** Schematic of the potential m6A sites in the *RRBP1* 3’-UTR. Shown below is the experimental design of constructing luciferase reporters containing WT or mutant 3’-UTR sequences of *RRBP1*. **H.** Luciferase reporter assay using the WT or mutated 3’-UTR constructs in DU145 cells with or without *METTL3*-KD. The firefly luciferase activity was normalized to Renilla luciferase activity. **I.** mRNA stability assay showing the kinetics of *RRBP1* expression in PCa cells with or without *METTL3*-KD after treatment with actinomycin D (10 µg/mL) for indicated time points. **J.** Efficient siRNA-mediated KD of RRBP1 both at the mRNA levels (left) and protein levels (right) in DU145 cells. Tubulin served as a loading control. **K** and **L.** Knocking down *RRBP1* inhibits clonal development (**K**) and migration (**L**) in DU145 cells. **M-O.** RRBP1 is a key downstream effector of METTL3/m6A pathway in PCa. Reducing the RRBP1 protein into baseline level (**M**) significantly counteracts the pro-proliferative (**N**) and pro-migratory (**O**) effects of METTL3 overexpression in PCa cells. Experiments were performed in METTL3 overexpressing DU145 cells treated with or without siRNA targeting *RRBP1.* Empty vector-overexpressing cells (OE-NC) transfected with siNC was used as a baseline control.

Next, we investigated the role of RRBP1 in PCa. siRNA-mediated KD of *RRBP1* significantly reduced its expression at both mRNA and protein levels (Fig. 5J), and concurrently suppressed cell proliferation (Fig. 5K) and migration (Fig. 5L). These results suggested an oncogenic role for RRBP1 in PCa progression. To further test whether RRBP1 was a key downstream effector of METTL3-mediated m6A pathway, we performed *RRBP1*-KD experiments in METTL3-overexpresssing PCa cells. Our data clearly showed that reducing the RRBP1 protein expression into a baseline level (comparable to cells without METTL3 overexpression; Fig. 5M) significantly counteracted the pro-proliferative (Fig. 5N) and pro-migratory (Fig. 5O) effects of METTL3 overexpression in DU145 cells. Altogether, our findings highlighted that METTL3 accelerates PCa progression via targeting *RRBP1* mRNA for stabilization in a m6A-dependent manner.

### 7. An m6A landscape in human PCa tissues reveals clinical significance of the METTL3/RRBP1 axis

A global m6A landscape in clinical PCa samples is lacking, although there are few studies that have identified multiple METTL3 targets at individual gene basis, with the aid of general m6A methylation patterns in particular PCa cell lines. Here, we for the first time performed m6A-seq on two pairs of matched cancer/paracancer tissues. By HOMER algorithm, we showed that transcripts pulled-down by an m6A-specific antibody markedly enriched for the known m6A consensus RRACH motif (Fig. 6A), validating our experimental pipeline. Moreover, m6A peaks were predominately located in the vicinity of 5’UTR to 1^st^ Exon and CDS to 3’UTR regions (Fig. 6B), similar to previously reported m6A maps generated in other tissues [32]. No obvious difference was found in the global m6A distribution pattern among cancer vs. paracancer tissues (Fig. 6C). Pair-wise comparison of m6A peaks in tumor vs. nontumor tissues identified 5180 methylated peaks in 2,910 genes and 3710 peaks in 2025 genes for patent 1 and 4, respectively (Fig. 6D and Table S4). To determine the cellular pathways that m6A might influence in clinical PCa, we conducted GO analysis on genes containing differential m6A peaks. Globally, m6A regulated mRNAs encoded a variety of pathways linked to a spectrum of important biological functions including development, cell adhesion and ECM (Fig. 6D), consistent with reported diverse roles of m6A pathway [2]. Specifically, besides that several GO terms were commonly enriched in two patients, a large part of enriched GO terms was still quite distinct, suggesting heterogeneity of m6A landscape among PCa patients. This is in line with the intrinsic property of heterogeneity of PCa. Our data therefore provides a useful resource for further mining m6A pattern in clinical PCa.

**Fig 6.**
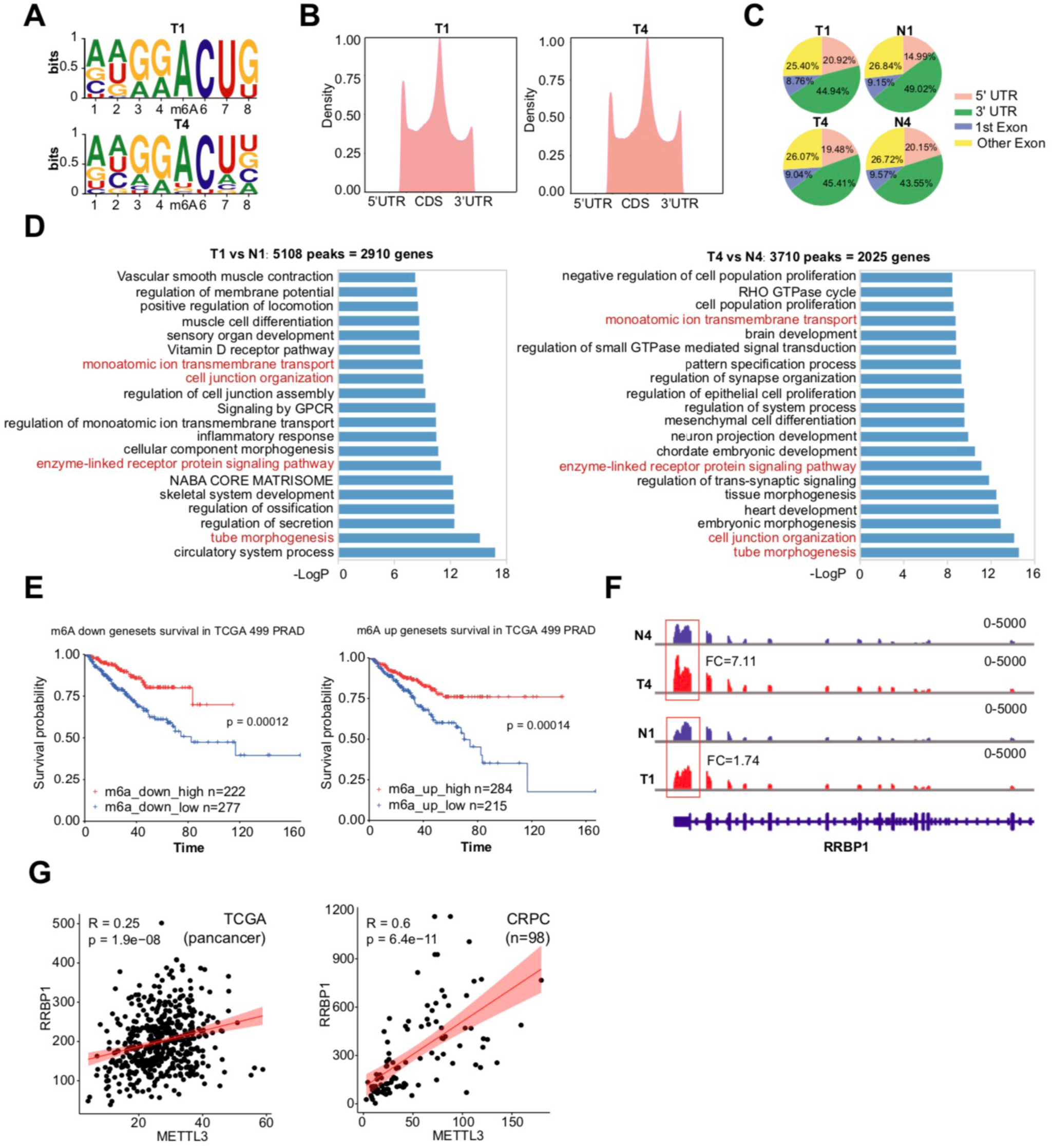
The clinical significance of the METTL3/RRBP1 axis in human PCa. **A.** Two paris of matched PCa and paracancer tissues were used for m6A-seq analysis. Shown are the concesus m6A RRACH motifs identified in two clinical prostate tumor samples. **B** and **C.** Statistics of m6A-seq peaks. Shown are the profiles of m6A peak density along mRNA transcript (**B**) and the genomic peak locations (**C**). **D.** GO analysis of genes bearing differential m6A peaks in tumor vs. nontumor tissues. Shown are the top 20 enriched pathways in two patient tumors, with a small proportion of GO pathways being commonly shared. **E.** Survival analysis of TCGA pan-cancer cohort based on genes that were co-upregulated or co-downregulated in both mRNA expression and m6A levels in tumor tissues. **F.** Distribution and differential enrichment of m6A peaks across *RRBP1* transcripts in tumor (T) and normal (N) samples. **G.** Positive correlations between *RRBP1* and *METTL3* mRNA levels in TCGA pri-PCa (left) and CRPC (right) cohorts.

To further demonstrate the clinical relevance of RNA m6A modification in PCa, we established signatures denoting to genes co-upregulated or co-downregulated in both mRNA expression and m6A levels in these two tumor tissues (Table S5). To our surprise, survival analysis of TCGA cohort showed that patients with higher expression of these co-regulated genes had a much worse prognosis (Fig. 6E). Considering that m6A methylation can both either promote or repress gene expression in an sustrate-dependent maner [3, 33], this data not only highlighted an involement of m6A modification in PCa biology but also pointed the complexcity of such regulatory mechnisam in regulating cancer behavior. Furthermore, in line with our cell line-derived data (Fig. 5G), we observed a greater enrichment of m6A siganls in *RRBP1* mRNA (mainly the 3’-UTR region) in tumor vs. non-tumor tissues (Fig. 6F). In addition, correlation analysis in both TCGA pri-PCa and CRPC datasets showed a strong positive correlation between *RRBP1* and *METTL3* at mRNA levels (Fig. 6G), further comfirming our *in vitro* experimental data that METTL3 upregulated *RRBP1* expression.

### 8. Peptide targeting METTL3-METTL14 interacting interface inhibites PCa *in vivo*

The protein-protein interaction (PPI) between METTL3 and METTL14, and the *S*-adenosyl methionine (AdoMet)-binding site of METTL3 are all crucial to the enzymatic activity of Writer complex [28]. With an aim to develop novel small-peptide METTL3 inhibitors, we designed two peptides corresponding to two interfaces of METTL3-METTL14 complex and one peptide to the AdoMet site of METTL3 (Fig. S4A), based on their structures elucidated previously [28]. Linking an arginine-9 motif (R9), a typical cationic and cell-penetrating peptide (CPP), to the N-terminal of these peptides made them potential inhibitors for m6A pathway (Fig. 7A). Using R9 as a negative control, we performed *in vitro* assays to show that peptides targeting interface 1 and 2 (i.e., R9-1 and R9-2) exhibited inhibitory effect on cell viability (Fig. 7B) and clonal development (Fig. 7C), with R9-2 being the superior one. Notably, peptide targeting the AdoMet site had no obvious impact on cell growth compared to R9 only (Fig. 7B and 7C). We therefore focused on R9-2 peptide for further investigation. Treatment of PCa cells with R9-2 significantly reduced the global m6A modification (Fig. 7D), which might be attributed to the reduced level of METTL3 protein after peptide treatment (Fig. 7E). Such reduction of METTL3 protein was consistent with the concept that a subunit is usually unstable if it is not properly assembled into a complex [34].

**Fig 7.**
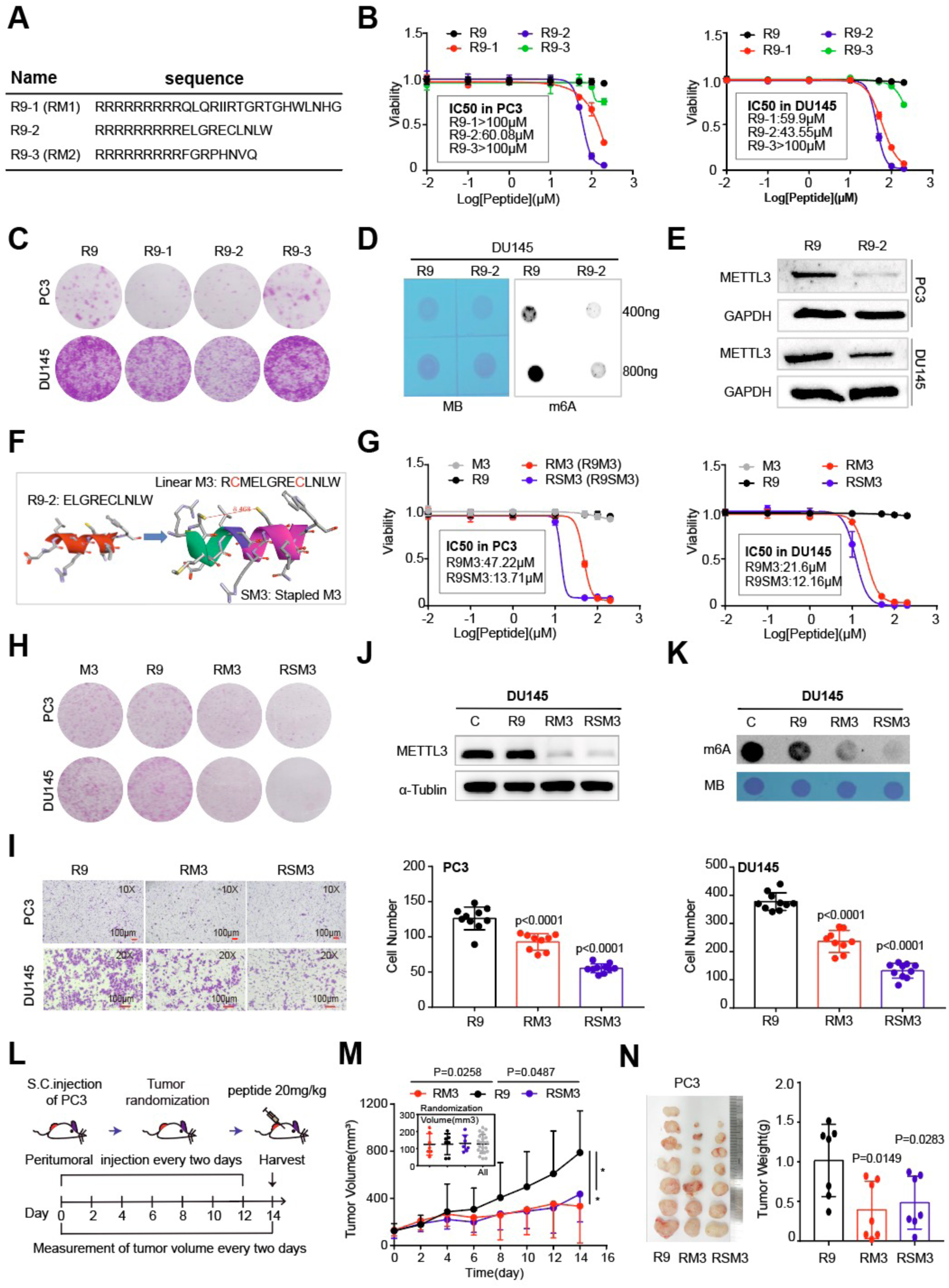
Small peptides targeting METTL3-METTL14 interaction inhibites PCa *in vitro* and *in vivo*. **A.** Sequence of peptides R9-1, R9-2, and R9-3. R9 denotes the cell penetrable arginine-9 motif. **B.** CCK8 cell viability (3 days) assays in indicated cells treated with different peptides at escalating doses. Data represent mean ± SD from a representative experiment with four technical repeats and the experiment was replicated three times with similar results. **C.** Colony formation assay in PC3 and DU145 cells treated with indicated peptides at IC50 dosages as revealed in B. **D** and **E.** Peptide R9-2 relative to control R9 treatment significantly reduces the global m6A levels (**D**) and METTL3 protein expression (**E**) in PC3 and DU145 cells. **F.** Schematic of devising a staple form of peptide R9-2. **G.** CCK8 cell viability (3 days) assays in indicated cells treated with different peptides at escalating doses. M3, linear form of the staple peptide. SM3, the stapled M3 peptide. **H** and **I.** PCa cells treated with RM3 and RSM3, compared to R9, peptides at IC50 markedly inhibits cell proliferation (**H**) and migration (**I**) *in vitro*. **J** and **K.** Reduced METTL3 protein expression (**J**) and the global m6A levels (**K**) in DU145 cells treated with RM3 and RSM3, compared to R9 or vehicle, peptides at IC50. The relative global m6A status was detected via dot blot assay. **L.** Schematic of *in vivo* peptide treatment. **M** and **N.** Inhibitory effects of METTL3-targeting peptides on the growth of PC3 CRPC model *in vivo*. Shown are the tumor growth curve (**M**; insets present tumor randomizations), endpoint tumor image and tumor weight (**N**) of xenografts (n = 7 for each group).

Although peptides were intrinsically safer than small molecule inhibitors, the IC50 of R9-2 in PC3 (60.08µM) and DU145 (43.55µM) cells were relatively high. Thus, we next sought to optimize it by modifying its sequence (i.e., increased 3 aa for the purpose of designing staple peptide) and changing it from linear form to a stapled form (Fig. 7F and S3A), which could increase peptide uptake and stability [35]. Using R9 or the linear sequence (before stapled; M3) as negative controls (which displayed no noticeable effect; Fig. 7G), we performed a series of cell-based assays to show that, relative to the linear RM3, stapled RSM3 displayed stronger inhibitory effect on cell proliferation (Fig. 7G) and clonal development (Fig. 7H) in two CRPC cell lines. As expected, RM3 or RSM3 treatment at IC50 concentration markedly reduced METTL3 protein abundance (Fig. 7J) and total m6A level (Fig. 7K). In migration assay, both R9M3 and R9SM3 halted cell mobility, with RSM3 being more effective than RM3 (Fig. 7I) in PCa cells. It is noteworthy that a first-in-class small-molecule METTL3 inhibitor STM2457 was recently discovered [6]. Albeit peptide vs. small molecule possessed intrinsically distinct properties, we compared the IC50s of STM2457 and our peptide inhibitors. STM2457 was initially described to be effective against multiple leukemia cell lines (IC50≈1.26 μM) [6], but we found that it was much less effective in killing solid PCa cells *in vitro*. The IC50s of STM2457 in both DU145 and PC3 cells were greater than 50 μM, whereas peptide inhibitors (RM3 and RSM3) comparatively behaved more superiorly (Fig. S4B), suggesting potential advantage of our peptide inhibitors over STM2457 in different biological context.

Last, we asked that whether an *in vitro* growth-inhibiting effect of METTL3-targeting peptides could be translated into therapeutic benefit *in vivo*. Using androgen-insensitive PC3 xenograft in immunodeficient mice as a CRPC model, we treated similar-sized tumors with indicated peptides at a dose of 20mg/Kg via peritumoral injection once every other day for 14 days (Fig. 7L). Results clearly showed that both R9M3 and R9SM3 alone effectively inhibited tumor growth in terms of tumor volume (Fig. 7M) and weight (Fig. 7N). Notably, no toxicity was noticed as evidenced by stable body weights during treatment course (Fig. S4C). Collectively, we have devised METTL3-targeting peptide inhibitors that may offer a new avenue for anticancer therapy.

## DISSCUSION

Compelling evidence has established METTL3 and/or m6A as critical players in variety of cancers in a context-dependent manner [3, 33], with majority of papers positioned METTL3 as an oncogene. Hence, METTL3 is widely pursued as a new target for anticancer drug development. Several reports have suggested an important role of METTL3 in PCa cell lines [14-18], with identification of a few downstream METTL3 targets on an individual gene basis. However, our understanding of m6A signaling in PCa is incomplete and a comprehensive and in-depth study of METTL3 is lacking. Here, we take an integrative and complementary approach to systematically dissect METTL3-mediated m6A pathway in PCa development and progression. Using both relatively indolent AR+ and aggressive AR- PCa models and human clinical samples, we perform *in vitro* and *in vivo* biological assays, mutagenesis, multi-omics sequencing, novel drug development, and preclinical therapeutic studies. ***First***, to directly show the clinical relevance of m6A modification, we perform m6A-seq in clinical PCa tumors to provide the actual m6A landscape, finding that m6A-modified transcripts are heterogeneously involved in many cancer-related pathways. ***Second***, METTL3 binds RNA to deposit m6A. We identify the global targets of METTL3 in PCa cells by RIP-seq technique. ***Third***, integrating the m6A-seq, RIP-seq, and regular RNA-seq data, we reveal a gene regulatory network centered on m6A signaling in aggressive PCa cells. ***Fourth***, using genetic KD and rescue experiments, we identify *RRBP1* as not only a direct target, but also a key functional downstream effector, of METTL3 in an m6A-dependent manner. A higher expression of *RRBP1* predicts a worse patient outcome. ***Fifth***, we develop novel small-peptide METTL3 inhibitors that demonstrate robust PCa-inhibiting activity *in vitro* and *in vivo*. These peptide inhibitors present a previously unexplored avenue for treating advanced PCa, especially CRPC.

Besides revealing the global m6A substrates in clinical tumor samples, we herein identify *RRBP1* as a new and direct substrate of METTL3 through multi-omics analysis. Mechanistically, we further show that upregulation of *RRBP1* in PCa is attributed, at least partially, to m6A-dependent stabilization of *RRBP1* mRNA catalyzed by METTL3. RRBP1 appears to be a key effector of METTL3’s oncogenic function in PCa, as *RRBP1*-KD abolishes the pro-proliferation impact caused by METTL3 overexpression. RRBP1 (reticulum ribosome-binding protein 1) is an endoplasmic reticulum (ER) membrane protein and is essential for ribosome binding and translocation of nascent proteins across the rough ER membrane [36-38]. Previous studies have linked an increased RRBP1 expression to pathology of various tumor types, including lung [39], breast [40], and bladder [41] cancers. These reports suggest RRBP1 as a potential oncogene, but little is known about its role in PCa. In this study, we experimentally show that depletion of RRBP1 significantly inhibits aggressive features of PCa cells, indicating that RRBP1 indeed functions oncogenically in PCa.

Another innovative and significant finding of our study is the development of small-peptide METTL3 inhibitors (i.e., linear RM3 and stapled RSM3). Based on the 3D structure of METTL3-METTL14 complex [28], we design multiple peptides and find that one targeting the interface 2 of PPI exhibits a good anticancer activity in PCa cells *in vitro*. Further sequence modification and stapling leads to the discovery of RSM3, which displays a superior toxicity against PCa both *in vitro* and *in vivo*. The design, detailed chemical characterization, and biological effect of such peptide inhibitor on other cancer types are presented in a parallel manuscript. While we are preparing the manuscript, a small-molecule METTL3 inhibitor STM2457 is reported to target blood cancers [6], whereas its application in solid tumors remains unstudied. Interestingly, our *in vitro* cell-killing assays show that the IC50 of STM2457 in PC3 and DU145 cells are both over 50 μM and 20 μM, respectively. Comparatively, the IC50 of peptide inhibitor RSM3 in PC3 and DU145 are 13.71µM and 12.11µM, respectively. Moreover, it is worth noting that although peptide (aiming to block METTL3-METTL14 interaction) and small molecule (aiming to block SAM domain) inhibitor function differently and possess distinct physicochemical properties, peptide drug is generally believed to deliver a higher selectivity and lower toxicity [35]. Our *in vivo* tumor assay also indicates RSM3 yields a better effect of restraining PC3 xenograft growth than STM2457 (see [parallel manuscript]). We also test the anticancer activity of RSM3 in some other cancer types such as pancreatic cancer [parallel manuscript], indicating a potential general application of METTL3 peptide inhibitor in many other METTL3-driven tumors. Collectively, our study reveals a novel METTL3/m6A/RRBP1 axis in aggressive PCa, which can be therapeutically targeted by small-peptide METTL3 antagonists.

## MATERIALS AND METHODS

### Cell culture

The human normal but transformed prostate epithelial cell line RWPE-1 and PCa cell lines PC3, DU145, LNCaP, and VCaP were purchased from Shanghai Cell Bank of the Chinese Academy of Sciences. RWPE-1 was cultured in K-SFM medium (Gibco, 17005-042). PCa cells were cultured in RPMI-1640 medium (Gibco, 8123209) supplemented with 10% heat-inactivated fetal bovine serum (FBS) (Excell, FSP500) and 1X antibiotics (penicillin/streptomycin) in humidified air at 37°C with 5% CO_2_. 293T cells were cultured in DMEM medium plus 8% FBS and 1X antibiotics (penicillin/streptomycin).

### RNA sequencing (RNA-seq)

The RNA-seq analysis was performed in collaboration with LC Sciences company. Briefly, total RNAs were isolated from METTL3-depleted or control PCa cells using Omega total RNA Midi Kit (Omega, R6834-02). Poly(A) RNA was subsequently purified used to generate cDNA libraries. All samples were sequenced on illumina Novaseq™ 6000 following the vendor’s recommended protocol. The clean reads were mapped to human reference genome (GRCh38) by STAR v2.7.3a [42]. Then, the DESeq2 package [43] was utilized to quantify gene expression and identified differentially expressed genes (DEGs) among different groups. Genes with absolute fold change (FC) ≥ 2, false discovery rate (FDR) < 0.05 and a baseMean (readcounts) >10 were identified as DEGs.

### RNA immunoprecipitation and high-throughput sequencing (RIP-seq)

The RIP experiment, high-throughput sequencing and data analysis were conducted by Seqhealth Technology Co., LTD (Wuhan, China). Briefly, PC3 and LNCaP cell lines were treated with cell lysis buffer. The 10% lysis sample was stored and named “input”, and 80% was used in immunoprecipitation reactions with anti-METTL3 antibody (Abcam, ab195352) and named “IP”, and 10% was incubated with rabbit IgG (Cell Signaling Technology) as a negative control and named “IgG”, respectively. Then the RNA of input and IP was extracted using TRIzol reagent (Invitrogen, USA). RNA was subsequently purified used to generate cDNA libraries. All samples were sequenced on illumina Novaseq™ 6000 with PE150 model. For data analysis, raw sequencing data was first filtered by Trimmomatic (version 0.36), low-quality reads were discarded and the reads contaminated with adaptor sequences were trimmed. Then, the clean reads were aligned to human reference genome (GRCh38) by STAR v2.7.3a [42]. The peak calling and annotation were conducted by exomePeak (Version 3.8) [44] and ChIPseeker [45] package.

### Methylated RNA Immunoprecipitation and sequencing (MeRIP-seq)

The MeRIP-seq (also called m6A-seq) is a method that aims to enrich for methylated RNA and ultimately identify modified transcripts. m6A-seq was performed in collaboration with LC Sciences company. Briefly, total RNA was isolated from prostate tissues using TriIzol reagent (Invitrogen, USA). Then, the poly(A)+ RNA was sheared into 100–200-nt fragments using Magnesium RNA Frag mentation Module (NEB, E6150S). A portion of poly(A)+ RNA fragment was directly used as input, and another portion of poly(A)+ RNA was used in immunoprecipitation reactions with anti-m6A antibody (Abcam, ab151230). Then the bound RNA was purified for downstream library preparation and sequencing on an illumina Novaseq™ 6000. For data analysis, the clean reads were aligned to human reference genome (GRCh38) by STAR v2.7.1a [42] and HOMER [46] was used for *de novo* and known motif identification. The Integrative Genomics Viewer (IGV) software was utilized to visualize selected peaks.

### MeRIP-RT-qPCR

To assess the m6A modification levels of a particular mRNA, m6A RIP was performed using the EpiQuik™ Cut&Run m6A RNA Enrichment (MeRIP) Kit (Epigentek, p9018) according to manufacturer’s instructions. Briefly, the isolated RNA was immunoprecipitated with m6A antibody-conjugated magnetic beads. The enriched RNA was then released, purified, and eluted. The relative m6A modification levels of RRBP1 were normalized to input. The primers for MeRIP RT-qPCR were shown as following: RRBP1-F: 5’-GCAGCCAGTTCTCAAAGG-3’ and RRBP1-R: 5’-AACACCCAGGAGGTCACA-3’.

### Plasmid construction and shRNA or siRNA transfection

To establish long-term knock-down (KD) cell lines, the Plko.1-puro-shRNA lentiviral vectors were constructed with the following two shRNAs: shMETTL3-718(5’-TGCACTTCAGACGAATTAT-3’) and shMETTL3-812(5’-TCAGTATCTTGGGCAAGTT-3’). To establish stable METTL3-overexpressing PCa cells, wild-type or catalytic-domain-mutated METTL3 (aa395-398, DPPW→APPA) was cloned into PCDH lentiviral vectors. Basic lentiviral procedures were previously described [47]. Lentivirus was produced in 293T packaging cells and titers determined using GFP positivity in 293T cells. PCa cells were infected with lentivirus, KD or overexpression efficiency was determined by Western blot at 72h post treatment. For siRNA-mediated KD experiments, the siRNA sequences were designed and synthesized by Gene Parma (Shanghai, China). The target sequences of siRNA were shown as following: METTL3-siRNA1 (5’-GCUACCUGGACGUCAGUAU-3’), METTL3-siRNA2 (5’-GGUUGGUGUCAAAGGAAAU-3’), and RRBP1-siRNA (5’-GCAUGUCGGUUA CAAGAAGAA-3’). When passaging, the PCa cells were plated in 6-well plates at a desired density and transfected with 50nM siRNA oligonucleotides. Transfection was performed with Lipo2000 in Opti-MEM medium (Gibico, Opti-MEM medium) and cells were subsequently used for experiments after 48h.

### Quantitative real-time PCR (qPCR)

RNA was isolated using RNA-easy Isolation Reagent (Vazyme, R701-01, China) following the manufacturer’s instructions. cDNA was generated by the Reverse Transcription Kit (Vazyme, R233-01). The qRT-PCR assay was performed using ChamQ SYBR Master Mix (Vazyme, Q311-02) on Bio-rad CFX96 real-time PCR system (Biorad, USA). All primers used in the present study were listed as following: GAPDH (F: 5’-ACTTTGGTATCGTGGAAGGACT-3’ and R: 5’-GCCTTGGCAGCGCCAGTAG-3’), METTL3 (F: 5’-GAGTGCATGAAAGCCAGTGA-3’ and R: 5’-ACTGGAATCACCTCCGACAC-3’), RRBP1 (F: 5’-AGTTCGGACCAGGTGAGGGAGCAC-3’ and R: 5’-GCGTCTTCAGCTGAACGGGGTC-3’).

### RIP-qPCR

The RIP assay was performed according to the previous description [48]. Briefly, the corresponding cell lysates were incubated with beads coated with 5 μg of control IgG antibody, anti-METTL3 antibody (Abcam, ab195352; 1:1000), with rotation at 4 °C overnight. Next, total RNA was extracted for the detection of RRBP1 expression by qRT-PCR. The primers for RIP RT-qPCR were shown as following: RRBP1-F: 5’-GCAGCCAGTTCTCAAAGG-3’ and RRBP1-R: 5’-AACACCCAGGAGGTCACA-3’.

### Luciferase report assay

The luciferase reporter assay was conducted as previously reported [49].DU145 cells were seeded in 24-well plates and transfected with pmirGLO luciferase reporter vector fused with RRBP1-3ʹUTR. The firefly luciferase and Renilla luciferase activity in each cell were detected by dual-luciferase reporter assay system (Promega, E1910) and the relative luciferase activity was further normalized to Renilla luciferase activity.

### RNA stability analyses

The RNA stability was determined according to a previous protoocol [50]. Briefly, PCa cells were cultured in 12-well plates overnight. Next, 10μg/ml actinomycin D (M4881, AbMole, USA) was added to cells to inhibit transcription for various time points as indicated. Then, RNA was extracted and determined by RT-qPCR. The RNA levels at different time points in indicated groups were calculated and normalized to GAPDH. The primers were the same as used for regular qPCR.

### Western blot

After washing with cold PBS, PCa cells were pelleted and resuspended in lysis buffer, incubated on ice with frequent vortex for 10 min. The supernatant was obtained by centrifugation at 12 000 g for 10min. Proteins were fractionated by SDSPAGE gel, transferred onto PVDF membranes, blocked in 5% nonfat milk or BSA in PBS/Tween-20, and then blotted with specific antibodies. Then, the membrane was washed, incubated with a secondary antibody, and washed again with wash buffer for three times. Finally, the membrane was exposed by Hyperfilm ECL kit and images were acquired. Antibodies against METTL3 (Abcam; ab195352), RRBP1 (Abclonal; A12239), GAPDH (Proteintech; 60004-1-Ig), α-Tubulin (Proteintech; 11224-1-AP) were all diluted at 1:1000 ratio.

### Cell proliferation and apoptosis assays

For cell proliferation assay, 2×10^4^ PCa cells were plated in 96-well plates and incubation at 37°C. The number of cells were counted after 72h, 120h and 168h. For colony formation assay, 5×10^3^ DU145, 8×10^3^ PC3, or 20×10^3^ LNCaP cells infected with lentivirus or siRNA were seeded in each well of 6-well plates and the medium was refreshed every 5 days. Colonies were fixed with methanol after 10 days for DU145 cells or 15 days for PC3 cells and then stained with 0.1% crystal violet (Solarbio, C8470) for 30 min followed by visualization. Cell viability was detected by adding 10% CCK8 (APExBIO, K1018, USA) into the cell cultures plated in 96-well plates and incubation at 37°C for 3 h at 72 h. The absorbance of each well was measured by a microplate reader set at 450 nm. All experiments were performed in triplicate. For cell apoptosis assay, Annexin V-PI apoptosis detection kit (Bioscience, A6030L) and Flow cytometry (Beckman, USA) were used according to the manufacturer’s instruction.

### Trans-well migration assay

Cell migration ability were measured by Trans-well assay. 5×10^4^ DU145 or 8×10^4^ PC3 cells suspended in 200 µL medium with 1% FBS were plated at the upper Trans-well chamber, and the entire chamber was placed into a 24-well plate with 20% FBS-containing medium placed at the bottom. After incubation for 1–2 days at 37°C, results were visualized by PROTOCOL™ Hema 3 staining kit (Fisher Scientific, Pittsburgh,741 PA). Images of the membranes were captured by Olympus IX71.

### Wound-healing assay

PCa cells in 6x-well culture plates were allowed to grow to 90% confluence, and a sterilized tip was utilized to introduce a scratch “wound” with the same width on the bottom of the dishes. We generally made two scratch wounds per well as technical replicates, and wound closure under multiple microscopic views per well was recorded. Images were captured at indicated time points to monitor the wound closing.

### RNA m6A dot-blot assay

The dot blot assay was performed according to a published bio-protocol paper (https://en.bio-protocol.org/e2095). Briefly, mRNA was isolated from total RNA and denatured and then spotted onto a Hybond-N+ membrane followed by crosslinking. Then, the membrane was washed and incubated first in blocking buffer and then with an anti-m6A antibody (Abcam, ab151230, USA) overnight at 4 °C with gentle shaking. The membrane was then washed, incubated with an anti-mouse antibody, and washed again. Finally, the membrane was exposed via Hyperfilm ECL kit and images were acquired. Methylene blue (MB) was used to stain mRNA and also as the loading control.

### Sphere-formation assays

For sphere-formation assays, cells were suspended in 1:1 Matrigel (BD Biosciences, San Jose, CA)/RPMI-1640 in a total volume of 100 μl. The mixtures were then plated around the rim of wells in a 12-well plate and allowed to solidify in 37 °C incubator for 25 min, followed by addition of 1ml of warm SC-medium (DMEM/F12 supplemented with 4μg/mL insulin, B27 (Invitrogen), and 20 ng/mL epidermal growth factor and basic fibroblast growth factor). Spheres that arose in 1–2 weeks with a diameter over 40 mm were counted. We usually used three escalating doses for each cell condition with 2-3 repeats for each cell dose.

### Animal experiments

Approximately 1 × 10^6^ PCa cells (PC3-shNC, PC3-sh812 cell lines) or 2× 10^6^ DU145 PCa cells (DU145-shNC, DU145-sh812, DU145-OENC, DU145-OEMETTL3-WT, DU145-OEMETTL3-Mut cell lines) per injection were suspended in a mixture of 100 μL PBS and Matrigel (1:1), and then subcutaneously injected into male BALB/c nude mice (6-7 weeks old). Tumor volume was monitored and calculated according to the equation of (length × width^2^)/2. Tumor weight were measured when mice were sacrificed.

### Peptide inhibitor treatment

Approximately 1×10^6^ PC3 cells were subcutaneously injected into male nude mice (n=21) to establish xenograft models. After 2 weeks post-injection, tumor-bearing mice were randomly assigned to three groups with a randomization rule of each group displaying similar tumor sizes. The three groups were separately treated with peptide R9, RM3, and RSM3 at a dosage of 20mg/Kg via orthotopic injection once every two days for 14 days (total 7 times). The size of engrafted tumors was measured at the interval of two days and tumor samples were collected at day 28 post the cancer cell implantation.

### Data availability

The RNA-seq, RIP-seq and m6A-seq data have been deposited in GEO database under accession code GSE241315, GSE240667, GSE240668, GSE240669, and GSE240672.

### Statistical analysis

We used the Log-Rank test to calculate P values for survival analyses. The co-expression analysis was performed by Spearman analysis. Otherwise, Student’s t-test, paired or unpaired two-tailed t-testwere used to calculate the statistical significance between comparisons depending on the data type. A p < 0.05 is considered statistically significant.

## Author contributions

D.Z. and J.S. conceived and designed the study, interpreted data, and finalized the manuscript. Y.F. and Z.L. performed most experiments. J.W. and C.Z. conducted Bioinformatic analysis. Y.T., J.X., Q.H., and W.L. provided assistance in *in vitro* cell-based assays. H.X. and B.X. provided reagents and clinical samples, respectively. All authors read and approved the manuscript.

