## Supplementary Figs. for "Stabilization of RRBP1 mRNA via an m6A-dependent manner in prostate cancer constitutes a therapeutic vulnerability amenable to small-peptide inhibition of METTL3"

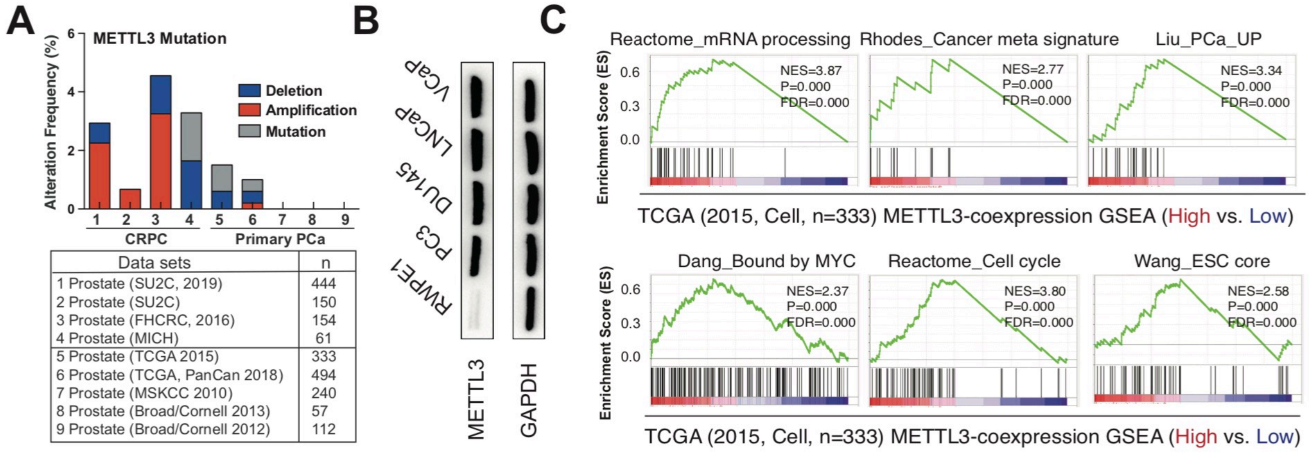


Figure S1

**Fig S1. METTL3 is overexpressed in human PCa cell lines and linked to multiple oncogenic biological pathways**

**A.** Bar plots illustrating the cumulative aberration frequencies of METTL3 across indicated cohorts.

**B.** Western blot analysis of METTL3 abundance in indicated prostatic cell lines. RWPE1 is an immortalized normal prostatic epithelial line. GAPDH served as a loading control.

**C.** Representative GSEA plots in genes correlated with *METTL3* expression in TCGA pri-PCa cohort.

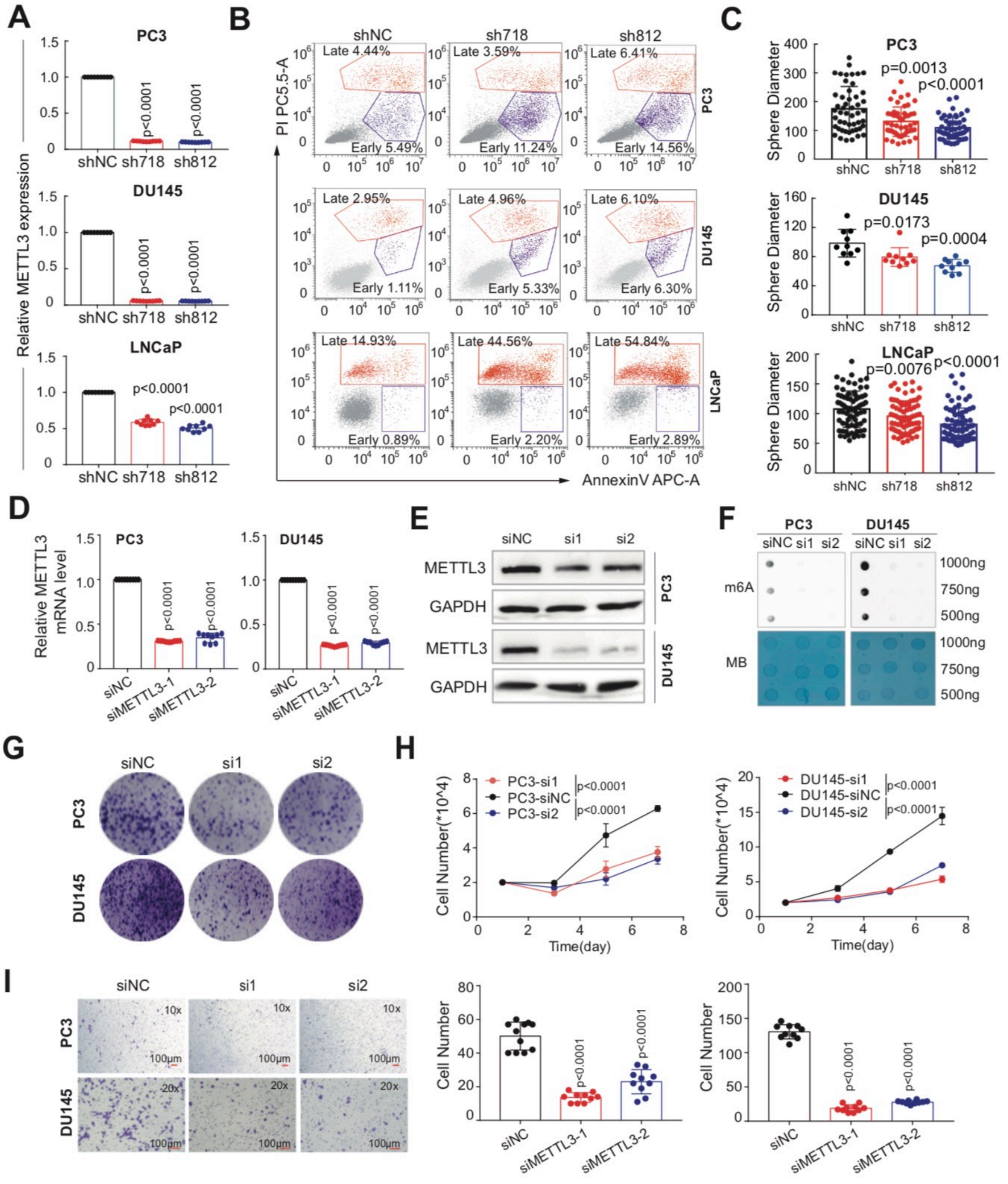

A
PC3
DU145
LNCaP
Relative METTL3 expression
shNC sh718 sh812
p<0.0001
B
shNC
sh718
sh812
PI PC5.5-A
AnnexinV APC-A
Late 4.44%
Early 5.49%
Late 3.59%
Early 11.24%
Late 6.41%
Early 14.56%
Late 2.95%
Early 1.11%
Late 4.96%
Early 5.33%
Late 6.10%
Early 6.30%
Late 14.93%
Early 0.89%
Late 44.56%
Early 2.20%
Late 54.84%
Early 2.89%
C
Sphere Diameter
p=0.0013
p<0.0001
p=0.0173
p=0.0004
p=0.0076
D
Relative METTL3 mRNA level
siNC siMETTL3-1 siMETTL3-2
E
siNC si1 si2
METTL3
GAPDH
F
m6A
MB
1000ng
750ng
500ng
G
H
PC3-si1
PC3-siNC
PC3-si2
DU145-si1
DU145-siNC
DU145-si2
Cell Number(*10^4)
Time(day)
I
10x
20x
100μm
Cell Number


Figure S2

**Fig S2. Biological role of METTL3 in human PCa cells**
**A.** qPCR analysis showing the knockdown efficiency of shRNAs targeting *METTL3* in indicated cell lines.
**B.** Depletion of *METTL3* significantly increases apoptosis measured by Annexin V staining assay.
**C.** Effect of *METTL3*-KD on sphere size in three PCa cell lines.
**D.** qPCR analysis showing the knockdown efficiency of siRNA-mediated KD of *METTL3* in PCa cells.
**E** and **F.** siRNA-mediated *METTL3*-KD significantly reduces METTL3 protein expression (**E**) and the global m6A levels (**F**) and in both PC3 and DU145 cells.
**G-I.** Knocking down *METTL3* by siRNAs inhibits clonal development (**G**), proliferation (**H**), and cell migration (**I**) in both PC3 and DU145 cells.
